# Aberrant cerebellar-default-mode functional connectivity underlying auditory verbal hallucinations in schizophrenia revealed by multi-voxel pattern analysis of resting-state functional connectivity MRI data

**DOI:** 10.1101/238519

**Authors:** Takashi Itahashi, Masaru Mimura, Sayaka Hasegawa, Masayuki Tani, Nobumasa Kato, Ryu-ichiro Hashimoto

## Abstract

Past neuroimaging studies have reported that aberrant functional connectivity (FC) underlying auditory verbal hallucinations (AVHs) in schizophrenia is highly distributed over multiple functional networks. There is thus a need for exploratory approaches without limiting analysis to particular seed regions or functional networks, to identify FC alterations underlying AVH. We applied a multi-voxel pattern analysis (MVPA) of FC together with a series of post-hoc FC analyses to resting-state fMRI data acquired from 25 patients with schizophrenia and 25 matched healthy controls. First, the MVPA revealed multiple clusters exhibiting altered FC patterns in schizophrenia. Subsequent multiple linear regression analysis using scores of these clusters identified that FC alteration in the right cerebellum crus I was significantly associated with the severity of AVH. Furthermore, post-hoc FC analysis with the right crus I as a seed revealed significant FC alterations with regions distributed across multiple functional networks, including speech, default-mode, thalamus, and cerebellum. Subsequent linear regression analyses further demonstrated that, among these regions, only reduced FC in the left precuneus was significantly associated with the severity of AVH. Our unbiased exploratory analysis of FC data revealed a novel evidence for the crucial role of FC between cerebellar and default-mode networks in AVH. (198 words)

## 1. Introduction

Auditory verbal hallucinations (AVH) are particularly common in patients with schizophrenia. AVH may occur in parallel with other symptoms, such as delusions, cognitive impairments, and social dysfunctions. In spite of intensive efforts using a broad range of technologies including functional imaging, the neural substrates of AVH have been highly elusive. Many past studies have primarily focused on abnormalities in auditory or receptive language areas associated with the experience of AVH. However, there is also abundant evidence for the involvement of non-language areas as well as homologous language areas in the non-dominant hemisphere in AVH. This has led to the proposal of various language and non-language-related hypotheses for AVH. For instance, the “inner-speech hypothesis” claims that AVH is generated by abnormal verbal imagery and action monitoring processes that are subserved in motor-related areas, including the supplementary motor area (SMA) (Moseley et al., 2013). The “cerebellar hypothesis” posits cerebellar dysfunctions as the core deficits underlying various schizophrenia symptoms including AVH (Andreasen and Pierson, 2008). Several other regions in limbic and subcortical structures, as well as cortical functional networks, may also deserve particular attention. These regions include the thalamus (Alonso-Solis et al., 2017; Shergill et al., 2000), hippocampus (Amad et al., 2014; Sommer et al., 2012), dorsal striatum (Rolland et al., 2015), sensorimotor areas (Wang et al., 2015), and default-mode network areas (Cui et al., 2017). These findings repeatedly suggest that critical abnormalities underlying AVH are not entirely confined to a particular brain region or functional network, but rather involve functional connectivity (FC) among brain regions dispersed across multiple functional systems.

There have been many studies regarding the relationships of altered FC with AVH and other schizophrenia symptoms by analyzing resting-state functional connectivity (RSFC) using various analytic methods (Berman et al., 2016; Yang et al., 2016; Yang et al., 2014; Zhu et al., 2016). Given the high degree of freedom in selecting the methodology, it is critical to select the optimal analytical methods according to the object of research. To date, seed-based analysis has probably been the most used method in RSFC studies. However, the selection of a small number of seed regions may lead to the risk of biasing interpretations of results, particularly when there are many suspected regions as in the case of AVH. On the other hand, independent component analysis (ICA) has been a highly powerful method for identifying functional networks without an a priori hypothesis regarding seed regions (McKeown et al., 2003). Although ICA is well-suited for detecting abnormal FC within a specific functional network, the regions suspected of involvement in AVH are dispersed across multiple functional networks, including auditory (Thoma et al., 2016), sensorimotor (Jardri et al., 2013), subcortical (Cui et al., 2017), and limbic (de la Iglesia-Vaya et al., 2014) systems. Therefore, the neural substrates of AVH may be more comprehensively identified by a novel analysis of FC that is more explorative and not restricted to particular seed regions or functional networks.

In this study, we adopted a data-driven approach that exploits principal component analysis (PCA) to characterize the inter-subject variability of FC patterns at the voxel-level in the brain (Amad et al., 2017; Whitfield-Gabrieli et al., 2015). This approach applied PCA to a set of FC maps from all participants with a given voxel as a seed. Individual differences in principal component (PC) scores indicate interindividual differences in patterns of FCs between that seed voxel and the rest of the brain. By repeating this step for all voxels followed by a group comparison of PC scores for every voxel in the brain, we aimed to identify clusters of voxels exhibiting altered FC patterns in schizophrenia (SZ) in an explorative data-driven manner. Because this analysis alone only indicates that such voxel clusters have atypical FCs with the rest of the brain, post-hoc FC analyses using those clusters as seeds are needed to identify specific FCs that are abnormal in SZ. Using this procedure, we aimed first to examine which brain regions, if any, exhibit atypical FC patterns associated with the severity of AVH in SZ. Post-hoc seed-based FC analyses were then used to identify specific FCs contributed to such atypical FC patterns. Owing to its highly data-driven nature, our study might reveal new critical inter-regional FCs contributing to AVH that have been unattended in the previous hypothesis-dependent approaches.

## 2. Materials and Methods

### 2.1. Participants

Twenty-five patients with SZ were recruited at the Karasuyama Hospital, Tokyo, Japan. Two experienced psychiatrists assessed all of the patients with SZ based on the Structured Clinical Interview for DSM-IV. No patients had other major psychiatric or neurological disease, severe somatic disorder, alcohol intake within 24 hours before the examination, or current or past alcohol or substance abuse. The psychiatric symptoms were assessed using the Positive and Negative Syndrome Scale (PANSS). In addition, the severity of auditory hallucination was evaluated using the Auditory Hallucination Rating Scale (AHRS) (Hoffman et al., 2003) and the hallucination item (P3) of PANSS. Although the P3 encompasses auditory as well as visual, olfactory, and cenesthetic hallucinations, we regarded the P3 as a measure of the severity of auditory hallucination. This is because patients in this study mainly experienced auditory hallucinations. The P3 score is referred to as the PANSS (H) score in this manuscript. Twenty-five matched healthy controls (HCs) participated in this study. Screening using a medical questionnaire ensured that no HC had any current or prior history of psychiatric illness. For each participant, handedness was assessed using the Edinburgh Handedness Inventory (Oldfield, 1971). The patient’s premorbid intelligence was estimated using the Japanese version of the National Adult Reading Test (Matsuoka et al., 2006). The same test was administered to the HCs to assess their putative intelligence.

Demographic and clinical data from the SZ and HC groups are summarized in Table 1. Age, gender, and handedness were matched between groups (all *p* > 0.9). This study was conducted in agreement with the principles of the Declaration of Helsinki and all of the procedures were approved by the Ethics Committee of the Faculty of Medicine of Showa University. After fully explaining the purpose of this study, written informed consent was obtained from all participants.

**Table 1:** The demographic and clinical data for the participants.

|  | HC | SZ | Statistics |  |
| --- | --- | --- | --- | --- |
| | Mean ± SD | Mean ± SD | *df* | *P*-value |
| Sample size (male/female) | 25 (15/10) | 25 (15/10) | - | - |
| Age (years) | 34.6 ± 8.20 | 34.4 ± 8.51 | 48 | 0.93 |
| Estimated IQ<sup>a</sup> | 106.86 ± 9.83 | 99.51 ± 9.45 | 48 | < 0.01 |
| Handedness (R/L) | 24/1 | 24/1 | 48 | 1.00 |
| Length of illness | - | 9.84 ± 7.16 | - | - |
| Chlorpromazine equivalents (mg/day) | - | 758 ± 515.91 | - | - |
| PANSS (P) score | - | 14.44 ± 5.75 | - | - |
| PANSS (H) score | - | 3.08 ± 1.80 | - | - |
| PANSS (N) score | - | 16.28 ± 5.19 | - | - |
| PANSS (G) score | - | 29.68 ± 7.59 | - | - |
| AHRS score | - | 14.6 ± 12.39 | - | - |

**Abbreviations:** HC: healthy control, SZ: schizophrenia, SD: standard deviation, IQ: intelligence quotient, PANSS: Positive and Negative Syndrome Scale, (P): positive scale, (H): hallucination item scale, (N): negative scale, (G): global psychopathology scale, and AHRS: Auditory Hallucination Rating Scale
<sup>a</sup>Estimated IQ was assessed by the Japanese version of the National Adult Reading Test (JART).

### 2.2. MRI acquisition

Functional and structural images were acquired using a 1.5 T GE Signa system (General Electric, Milwaukee, WI, USA) with a phased-array whole-head coil. Functional images were acquired using a gradient echo-planar imaging sequence (in-plane resolution: 3.4375 × 3.4375 mm, echo time [TE]: 40 ms, repetition time [TR]: 2,000 ms, flip angle: 90°, slice thickness: 4 mm with a 1 mm slice gap, matrix size: 64 × 64, 27 axial slices). Two-hundred and eight volumes were acquired in a single run. The first four volumes were discarded to allow for T1 equilibration. During the resting-state scan, the participant was instructed to lie relaxed inside the scanner with his/her eyes closed, and yet to stay awake in the dim scanner room. A T1-weighted spoiled gradient recalled three-dimensional MRI image was acquired (in-plane resolution: 0.9375 × 0.9375 mm, 1.4 mm slice thickness, TR: 25 ms, TE: 9.2 ms, matrix size: 256 × 256, 128 sagittal slices) for normalization purposes.

### 2.3 rs-fMRI data preprocessing

The collected data were preprocessed using Statistical Parametric Mapping software (SPM8; Wellcome Department of Cognitive Neurology, London, UK). The data underwent a series of preprocessing steps in the following order: (1) slice-timing correction, (2) head motion correction, (3) normalization and resampling to a resolution of 2 × 2 × 2 mm, and (4) spatial smoothing using a Gaussian kernel (6mm full-width at half-maximum).

Nuisance signals, including six head motion parameters, their temporal derivatives, mean signals from white matter, and mean signals from cerebrospinal fluid, were regressed out from the signals in each voxel. Of note, global signal regression was not performed since its validity is still unclear (Murphy et al., 2009; Saad et al., 2012). A band-pass filter (0.009-0.08 Hz) was applied to reduce the effects of low-frequency drifts and physiological noises. To reduce spurious changes in FC due to subtle head motions, we calculated a frame-wise displacement (FD) and removed volumes with FD > 0.5 mm by employing the scrubbing method (Power et al., 2012). There was no significant between-group difference in the mean FD (HC: 0.11 ± 0.04 [mean ± standard deviation]; SZ: 0.12 ± 0.04; *p* = 0.68, *t* = -0.41).

### 2.4. Functional connectivity analyses

#### 2.4.1. Multi-voxel pattern analysis for FC

To identify altered FC patterns associated with SZ and its core symptom of AVH, we adopted an exploratory data-driven approach, which is known as a multi-voxel pattern analysis (MVPA) of FC, as described by Whitfield-Gabrieli et al. (2015). Figure 1 shows an overview of the method. Briefly, we first generated a study-specific brain mask by including only voxels present in all 50 participants and those survived following the application of a 25% gray matter probability mask. For each participant, we regarded each voxel in the mask as a seed and calculated FC of the voxel with the rest of voxels in the mask. This procedure resulted in a data matrix for each seed, each row of which corresponded to a seed-to-voxel FC pattern of a single participant. PCA reduced the dimensionality of this matrix while preserving the explained inter-subject variability in the matrix by a lower number of PCs. We used this procedure to generate PC score maps for each participant. Among the resulting PC score maps, the 1st PC score map was selected as an input to the second-level analysis for the group comparison between HCs and patients with SZ, because the 1st PC score explains the largest portion of the inter-subject variability in the data. For the sake of simplicity, we will refer to the 1st PC score map as the PC map hereafter.

**Figure 1:**
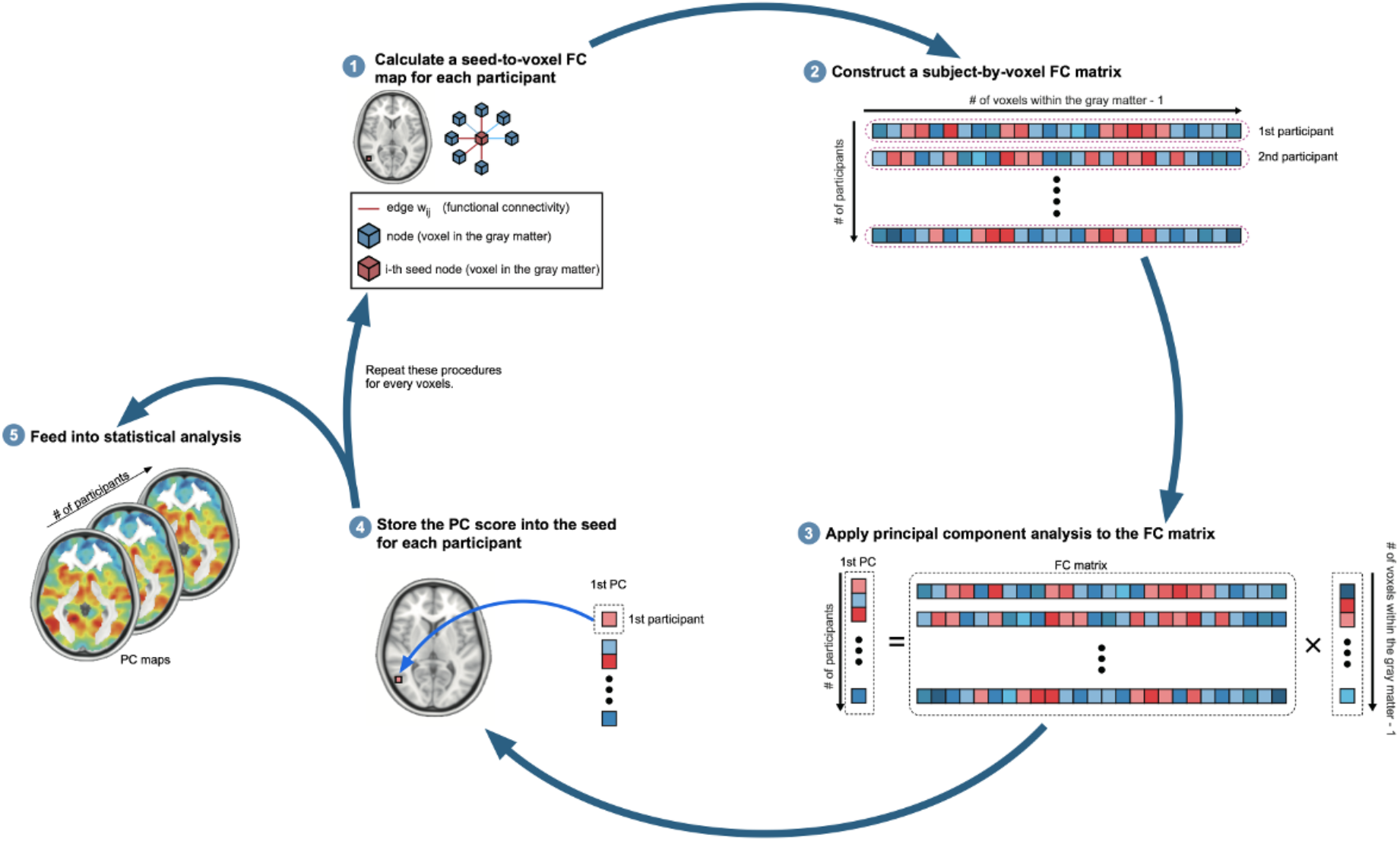
Overview of multi-voxel pattern analysis of functional connectivity. First, we regarded a single voxel as a seed region and then calculated functional connectivity (FC) between the seed and the rest of the voxels in the brain. Second, we constructed a subject-by-voxel FC matrix and applied principal component analysis (PCA) to this matrix. Third, we assigned the 1st principal component (PC) score to the seed voxel. We repeated this procedure for all voxels to generate a PC map for each subject. Finally, we performed group comparisons on these PC maps.

For group comparisons, two-sample *t*-tests were applied to the set of the PC maps, while age and the mean FD were included as nuisance covariates. Since we examined both directions separately (i.e., SZ < HC and SZ > HC) for each voxel, the statistical threshold was set to 0.025 after family-wise error (FWE) correction. The spatial extent threshold of 20 voxels was further applied to identify significant clusters.

The second-level analysis identified multiple (i.e., 25) brain regions showing significant group differences. From among these regions, we then identified regions associated with the severity of AVH by adopting a stepwise linear regression analysis. We first calculated the averaged PC score for each brain region and then used the series of 25 mean PC scores as independent variables for predicting either of the two clinical measures of AVH [i.e., AHRS or PANSS (H)]. When step-wise linear regression detected such associations for any region, we calculated a Spearman’s rank correlation coefficient between the PC score of that region and the clinical score. The statistical threshold was set to 0.05 for stepwise regression and correlation analyses.

#### 2.4.2. Post-hoc seed-based FC analysis

Regions with significant group differences in PC scores observed in the MVPA of FC had different FC patterns with some regions in the rest of the brain in patients with SZ. To identify the specific FCs that contributed to atypical PC scores in the SZ group, we performed post hoc seed-based analysis on a cluster showing significant between-group differences. To avoid repeating group comparisons multiple times, we focused only on the right crus I, because only this region consistently exhibited significant between-group difference and a significant association with the severity of AVH as measured by the PANSS (H) score.

For group comparisons of the seed-based FC maps, two sample *t*-tests were performed on each voxel while including age and the mean FD as nuisance covariates. The voxel-based statistical threshold was set to 0.01 after false discovery rate (FDR) correction. The spatial extent threshold of 20 voxels was also used to identify significant clusters.

Post-hoc seed-based analysis identified multiple brain regions exhibiting significant between-group differences. To further explore the specific FCs with the right crus I that were associated with the severity of AVH, we performed multiple linear regression analysis using either PANSS (H) or AHRS on the right crus I-seeded FC maps within the SZ group. Before the analysis, we generated a mask to restrict statistical tests to only voxels with significant between-group differences. The statistical threshold was set to 0.001 at the voxel level. The extent threshold was estimated using 3dFWHMx and 3dClustsim implemented in Analysis of Functional Neuroimages (AFNI; http://afni.nimh.nih.gov/afni). The threshold was set to 47 voxels to achieve an FWE rate of *p* < 0.05 at the cluster level (Cox, 1996). The above analysis included age, handedness, gender, and the mean FD as nuisance covariates.

## 3. Results

### 3.1. MVPA of FC

Group comparisons of PC maps revealed multiple clusters with significantly altered PC scores in the SZ group when compared to the HC group (Figure 2 and Table 2). The SZ group had reduced scores in the left superior temporal gyrus (STG), thalamus, hippocampus, bilateral lingual gyri (LING), and crus I. Patients with SZ also had increased scores in the left precentral gyrus/postcentral gyrus (PreCG/PostCG), right SMA, bilateral thalamus, LING, and crus I. These results indicate that there are significantly altered FCs involving these regions.

**Table 2:**
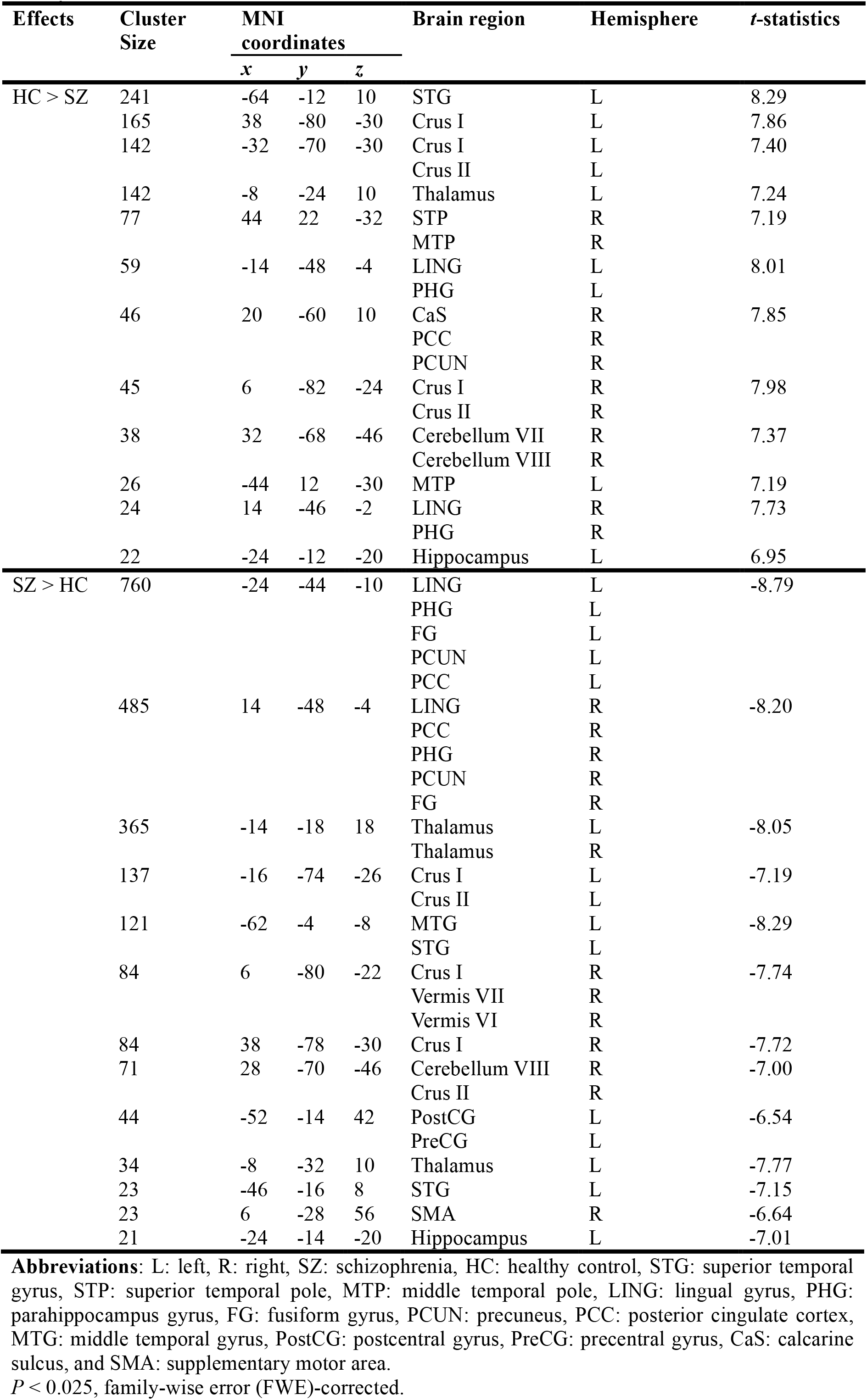
Alterations of 1st principal component score in patients with schizophrenia relative to healthy controls.

| Effects | Cluster Size | MNI coordinates |  |  | Brain region | Hemisphere | *t*-statistics |
| --- | --- | --- | --- | --- | --- | --- | --- |
|  |  | *x* | *y* | *z* |  |  |  |
| HC > SZ | 241 | -64 | -12 | 10 | STG | L | 8.29 |
|  | 165 | 38 | -80 | -30 | Crus I | L | 7.86 |
|  | 142 | -32 | -70 | -30 | Crus I | L | 7.40 |
|  |  |  |  |  | Crus II | L |  |
|  | 142 | -8 | -24 | 10 | Thalamus | L | 7.24 |
|  | 77 | 44 | 22 | -32 | STP | R | 7.19 |
|  |  |  |  |  | MTP | R |  |
|  | 59 | -14 | -48 | -4 | LING | L | 8.01 |
|  |  |  |  |  | PHG | L |  |
|  | 46 | 20 | -60 | 10 | CaS | R | 7.85 |
|  |  |  |  |  | PCC | R |  |
|  |  |  |  |  | PCUN | R |  |
|  | 45 | 6 | -82 | -24 | Crus I | R | 7.98 |
|  |  |  |  |  | Crus II | R |  |
|  | 38 | 32 | -68 | -46 | Cerebellum VII | R | 7.37 |
|  |  |  |  |  | Cerebellum VIII | R |  |
|  | 26 | -44 | 12 | -30 | MTP | L | 7.19 |
|  | 24 | 14 | -46 | -2 | LING | R | 7.73 |
|  |  |  |  |  | PHG | R |  |
|  | 22 | -24 | -12 | -20 | Hippocampus | L | 6.95 |
| SZ > HC | 760 | -24 | -44 | -10 | LING | L | -8.79 |

|  |  |  |  |  |  |  |
| --- | --- | --- | --- | --- | --- | --- |
|  |  |  |  | PHG | L |  |
|  |  |  |  | FG | L |  |
|  |  |  |  | PCUN | L |  |
|  |  |  |  | PCC | L |  |
| 485 | 14 | -48 | -4 | LING | R | -8.20 |
|  |  |  |  | PCC | R |  |
|  |  |  |  | PHG | R |  |
|  |  |  |  | PCUN | R |  |
|  |  |  |  | FG | R |  |
| 365 | -14 | -18 | 18 | Thalamus | L | -8.05 |
|  |  |  |  | Thalamus | R |  |
| 137 | -16 | -74 | -26 | Crus I | L | -7.19 |
|  |  |  |  | Crus II | L |  |
| 121 | -62 | -4 | -8 | MTG | L | -8.29 |
|  |  |  |  | STG | L |  |
| 84 | 6 | -80 | -22 | Crus I | R | -7.74 |
|  |  |  |  | Vermis VII | R |  |
|  |  |  |  | Vermis VI | R |  |
| 84 | 38 | -78 | -30 | Crus I | R | -7.72 |
| 71 | 28 | -70 | -46 | Cerebellum VIII | R | -7.00 |
|  |  |  |  | Crus II | R |  |
| 44 | -52 | -14 | 42 | PostCG | L | -6.54 |
|  |  |  |  | PreCG | L |  |
| 34 | -8 | -32 | 10 | Thalamus | L | -7.77 |
| 23 | -46 | -16 | 8 | STG | L | -7.15 |
| 23 | 6 | -28 | 56 | SMA | R | -6.64 |
| 21 | -24 | -14 | -20 | Hippocampus | L | -7.01 |

**Abbreviations:** L: left, R: right, SZ: schizophrenia, HC: healthy control, STG: superior temporal gyrus, STP: superior temporal pole, MTP: middle temporal pole, LING: lingual gyrus, PHG: parahippocampus gyrus, FG: fusiform gyrus, PCUN: precuneus, PCC: posterior cingulate cortex, MTG: middle temporal gyrus, PostCG: postcentral gyrus, PreCG: precentral gyrus, CaS: calcarine sulcus, and SMA: supplementary motor area.
$P < 0.025$ , family-wise error (FWE)-corrected.

**Table 3:**
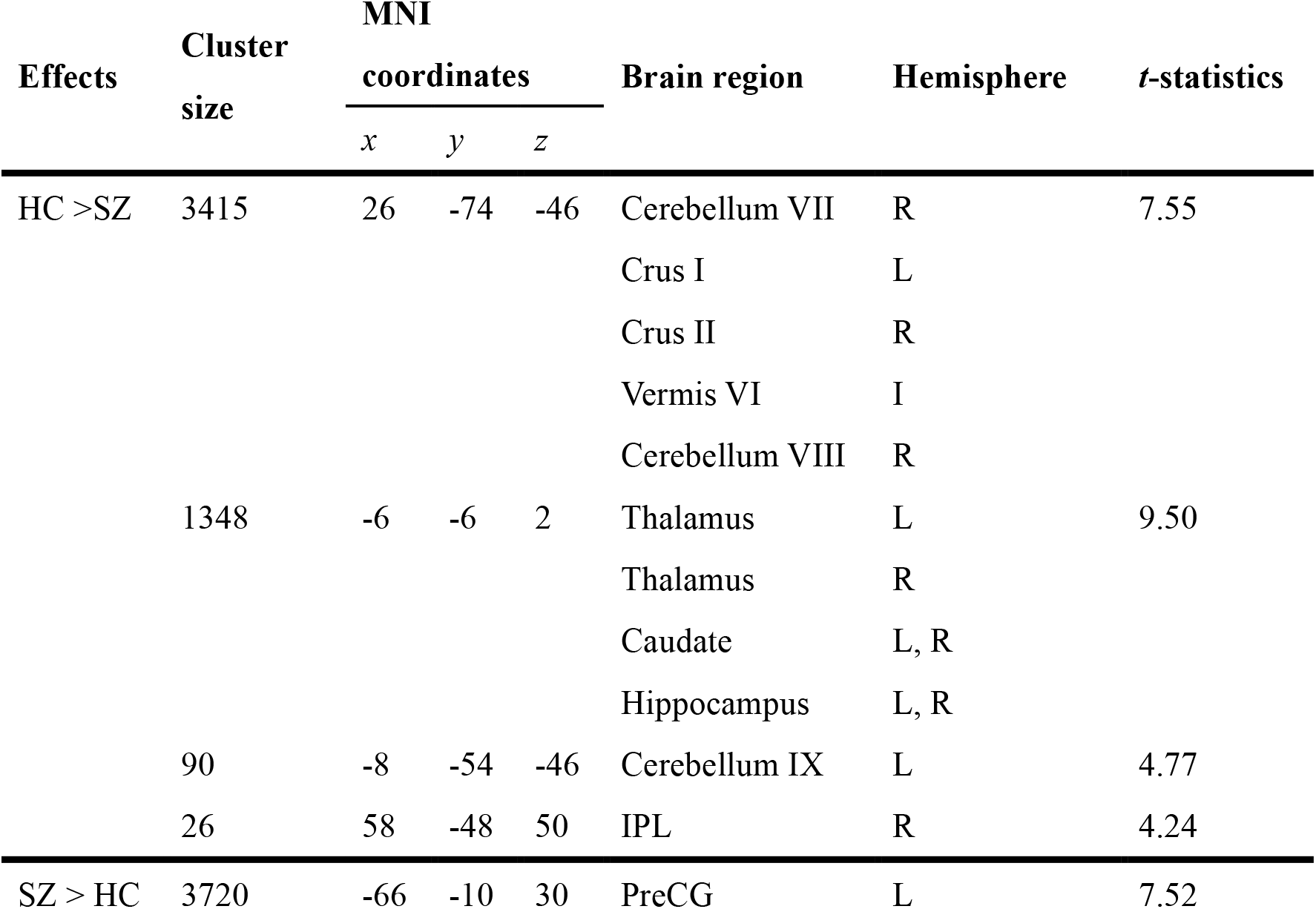
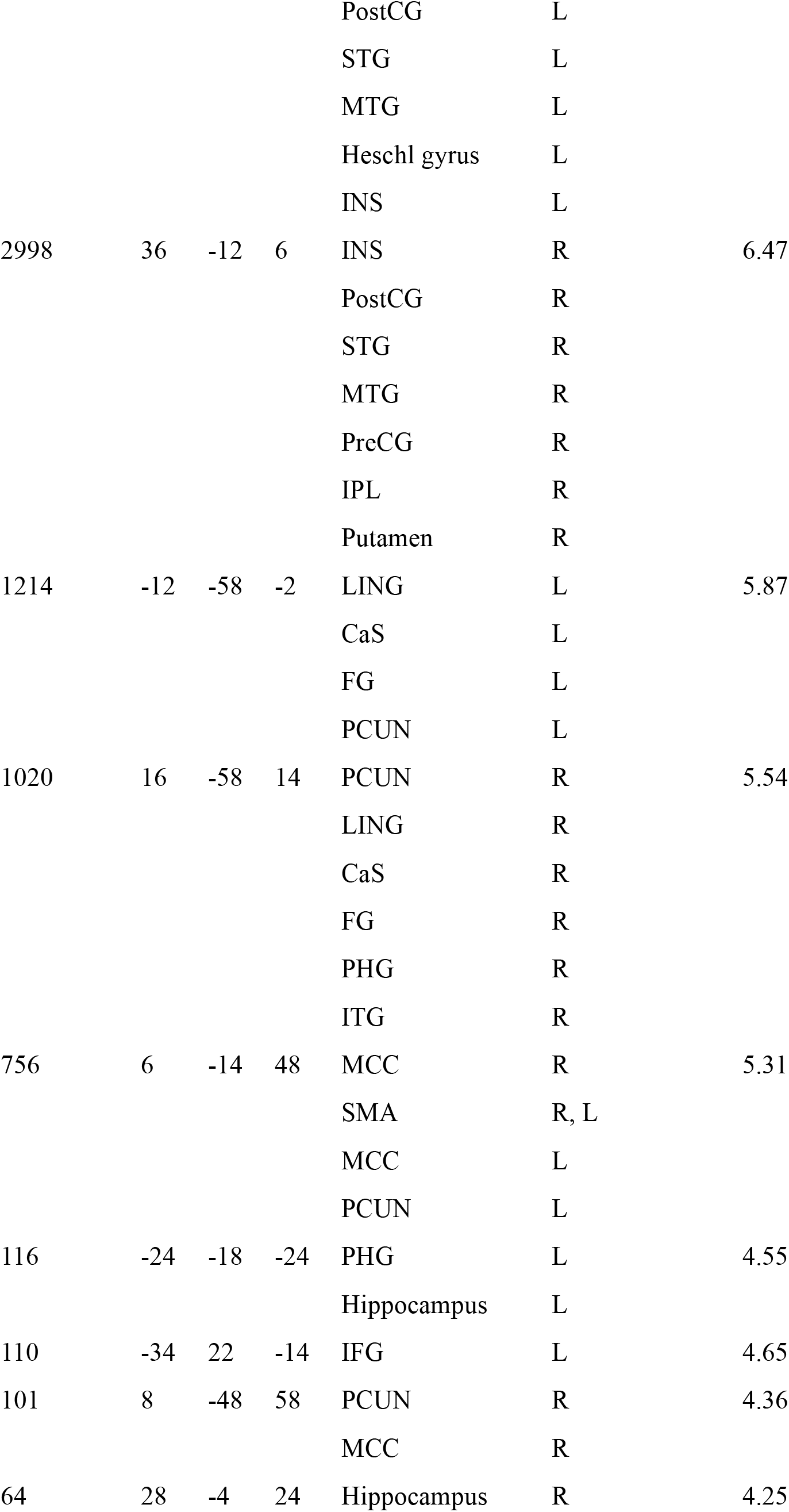
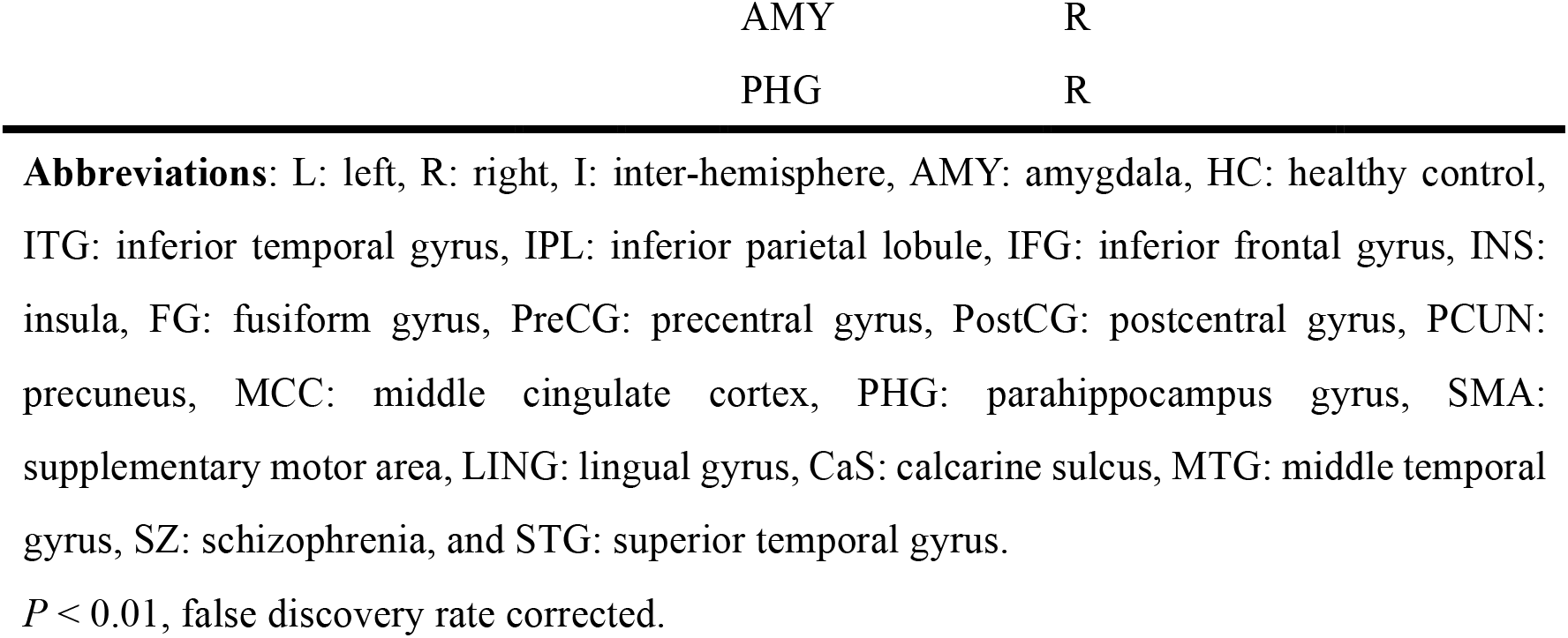
Altered right crus I seed-based functional connectivity in patients with schizophrenia when compared to healthy controls.

| Effects | Cluster size | MNI |  |  | Brain region | Hemisphere | *t*-statistics |
| --- | --- | --- | --- | --- | --- | --- | --- |
|  |  | coordinates |  |  |  |  |  |
|  |  | *x* | *y* | *z* |  |  |  |
| HC > SZ | 3415 | 26 | -74 | -46 | Cerebellum VII | R | 7.55 |
|  |  |  |  |  | Crus I | L |  |
|  |  |  |  |  | Crus II | R |  |
|  |  |  |  |  | Vermis VI | I |  |
|  |  |  |  |  | Cerebellum VIII | R |  |
|  | 1348 | -6 | -6 | 2 | Thalamus | L | 9.50 |
|  |  |  |  |  | Thalamus | R |  |
|  |  |  |  |  | Caudate | L, R |  |
|  |  |  |  |  | Hippocampus | L, R |  |
|  | 90 | -8 | -54 | -46 | Cerebellum IX | L | 4.77 |
|  | 26 | 58 | -48 | 50 | IPL | R | 4.24 |
| SZ > HC | 3720 | -66 | -10 | 30 | PreCG | L | 7.52 |

|  |  |  |  |  |  |  |
| --- | --- | --- | --- | --- | --- | --- |
|  |  |  |  | PostCG | L |  |
|  |  |  |  | STG | L |  |
|  |  |  |  | MTG | L |  |
|  |  |  |  | Heschl gyrus | L |  |
|  |  |  |  | INS | L |  |
| 2998 | 36 | -12 | 6 | INS | R | 6.47 |
|  |  |  |  | PostCG | R |  |
|  |  |  |  | STG | R |  |
|  |  |  |  | MTG | R |  |
|  |  |  |  | PreCG | R |  |
|  |  |  |  | IPL | R |  |
|  |  |  |  | Putamen | R |  |
| 1214 | -12 | -58 | -2 | LING | L | 5.87 |
|  |  |  |  | CaS | L |  |
|  |  |  |  | FG | L |  |
|  |  |  |  | PCUN | L |  |
| 1020 | 16 | -58 | 14 | PCUN | R | 5.54 |
|  |  |  |  | LING | R |  |
|  |  |  |  | CaS | R |  |
|  |  |  |  | FG | R |  |
|  |  |  |  | PHG | R |  |
|  |  |  |  | ITG | R |  |
| 756 | 6 | -14 | 48 | MCC | R | 5.31 |
|  |  |  |  | SMA | R, L |  |
|  |  |  |  | MCC | L |  |
|  |  |  |  | PCUN | L |  |
| 116 | -24 | -18 | -24 | PHG | L | 4.55 |
|  |  |  |  | Hippocampus | L |  |
| 110 | -34 | 22 | -14 | IFG | L | 4.65 |
| 101 | 8 | -48 | 58 | PCUN | R | 4.36 |
|  |  |  |  | MCC | R |  |
| 64 | 28 | -4 | 24 | Hippocampus | R | 4.25 |

|  |  |
| --- | --- |
| AMY | R |
| PHG | R |

**Abbreviations:** L: left, R: right, I: inter-hemisphere, AMY: amygdala, HC: healthy control, ITG: inferior temporal gyrus, IPL: inferior parietal lobule, IFG: inferior frontal gyrus, INS: insula, FG: fusiform gyrus, PreCG: precentral gyrus, PostCG: postcentral gyrus, PCUN: precuneus, MCC: middle cingulate cortex, PHG: parahippocampus gyrus, SMA: supplementary motor area, LING: lingual gyrus, CaS: calcarine sulcus, MTG: middle temporal gyrus, SZ: schizophrenia, and STG: superior temporal gyrus.
$P < 0.01$ , false discovery rate corrected.

**Figure 2:**
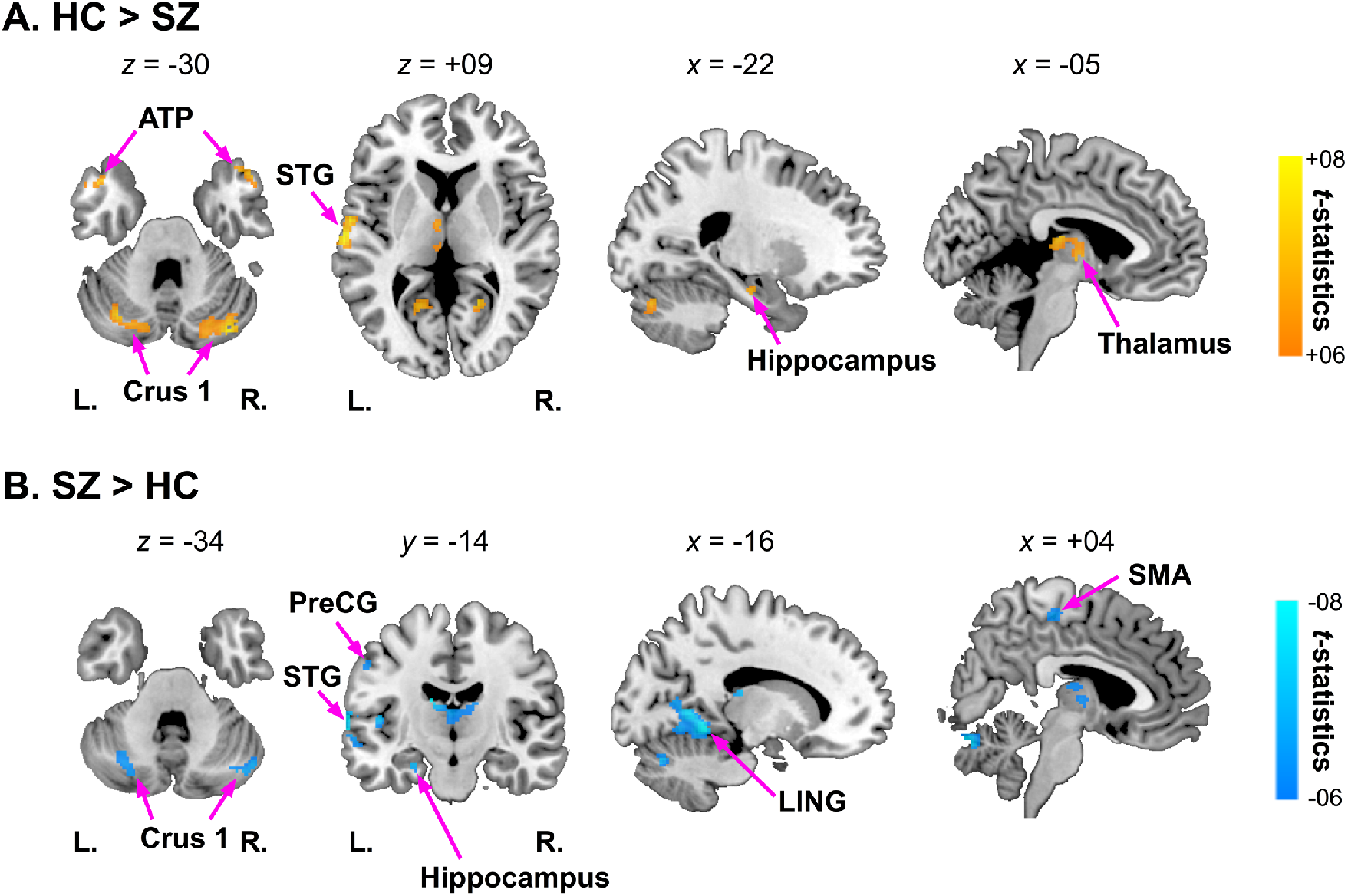
Altered principal component score of functional connectivity pattern in patients with schizophrenia when compared to health controls. The *t*-statistics of the group comparisons of the 1st principal component score were shown (health control [HC] > schizophrenia [SZ] and SZ > HC). Multiple comparisons were corrected based on the family-wise error correction (significance level: *p* < 0.025, corrected), and an extent threshold of 20 voxels was further applied. Orange-yellow colors represent positive *t*-statistics, which indicate significant reductions in the SZ group when compared to the HC group. Light-blue colors represent negative t-statistics, which indicate significant increases in the SZ group when compared to the HC group. R: right, L: left, ATP: anterior temporal pole, LING: lingual gyrus, SMA: supplementary motor area, PreCG:precentral gyrus, and STG: superior temporal gyrus.

We examined the specific brain regions that contribute to the severity of AVH from among the multiple brain regions identified in the MVPA of FC. We performed a stepwise linear regression analysis using the mean PC scores of regions with significant group differences as regressors for either the PANSS (H) score or AHRS. This analysis revealed a significant association between altered PC scores in the right crus I (peak MNI coordinate = [+38, -78, -30]) and the PANSS (H) scores (*p* = 0.0278). In this region, association with the AHRS also approached the significance level (*p* = 0.058). The spearman’s rank correlation coefficient also confirmed a significant positive correlation between altered PC scores in the right crus I and the PANSS (H) scores (Spearman’s *r* = 0.46, *p* = 0.021). A tendency for a positive correlation was also found with the AHRS (Spearman’s *r* = 0.38, *p* = 0.058) (Figure 3).

**Figure 3:**
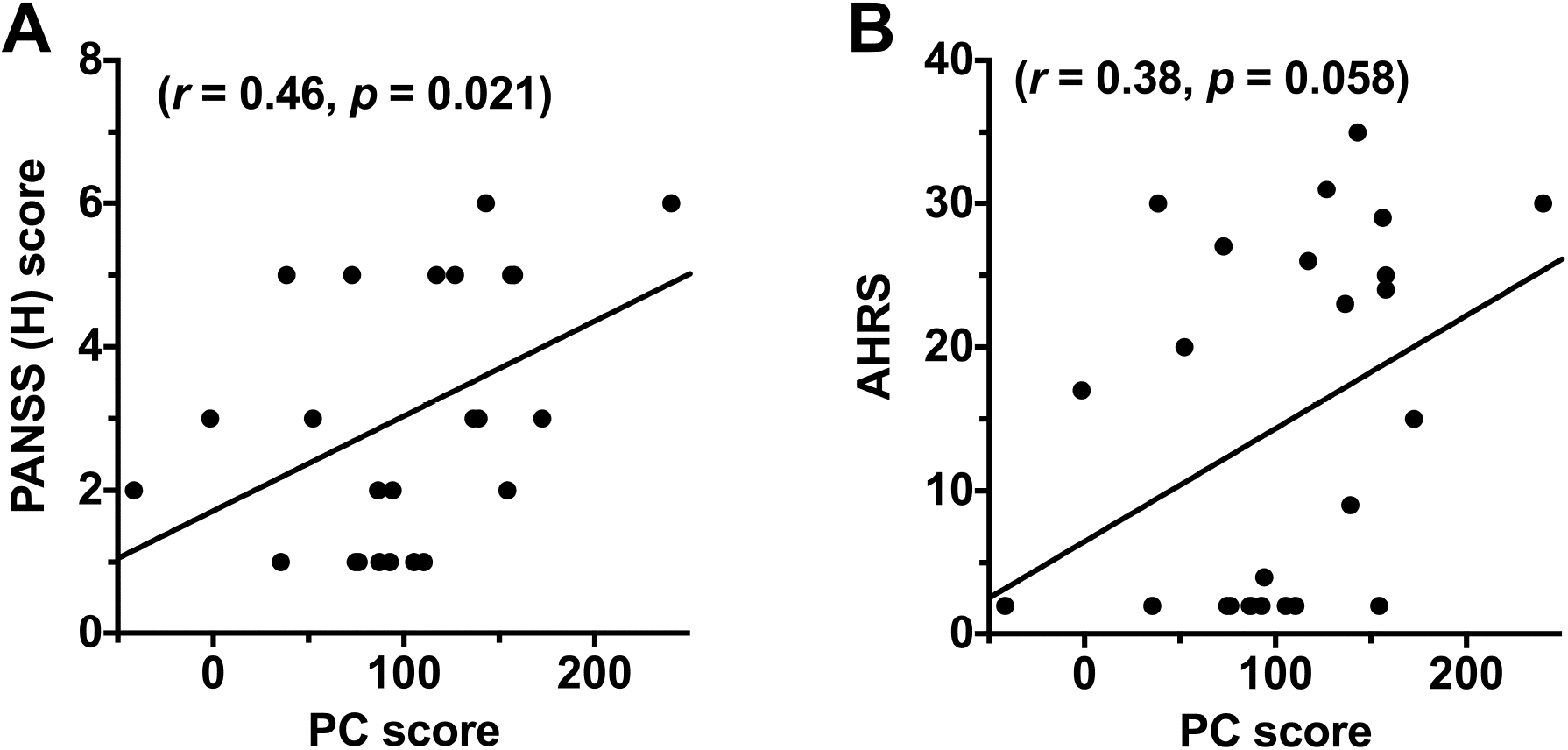
Positive correlations between altered principal component scores and the severity of auditory verbal hallucinations. Stepwise linear regression identified that altered principal component (PC) scores in the right crus I was significantly contributed to the prediction of the severity of auditory verbal hallucinations (AVHs) as assessed using the Positive and Negative Syndrome Scale (PANSS) (H) score. To confirm the association between altered PC score in this region and PANSS (H) score, correlation analysis was performed using Spearman’s rank correlation coefficient. (A) Scatterplot of association between altered PC score in the right crus I and the PANSS (H) score. (B) Scatterplot of association between altered PC score in the right crus I and the Auditory Hallucination Rating Scale.

### 3.2. Post-hoc seed-based FC analysis

Because the MVPA does not specifically identify FCs that contributed to altered PC scores, we performed a post-hoc seed-based analysis with the identified region (right crus I) as a seed. We found that the SZ group exhibited hyper-connectivity mainly with brain regions involved in speech perception and production, such as the bilateral STG, bilateral PreCG, and bilateral insula. Regions with hypo-connectivity were mainly observed in the cerebellum, thalamus, and caudate nucleus (Figure 4).

**Figure 4:**
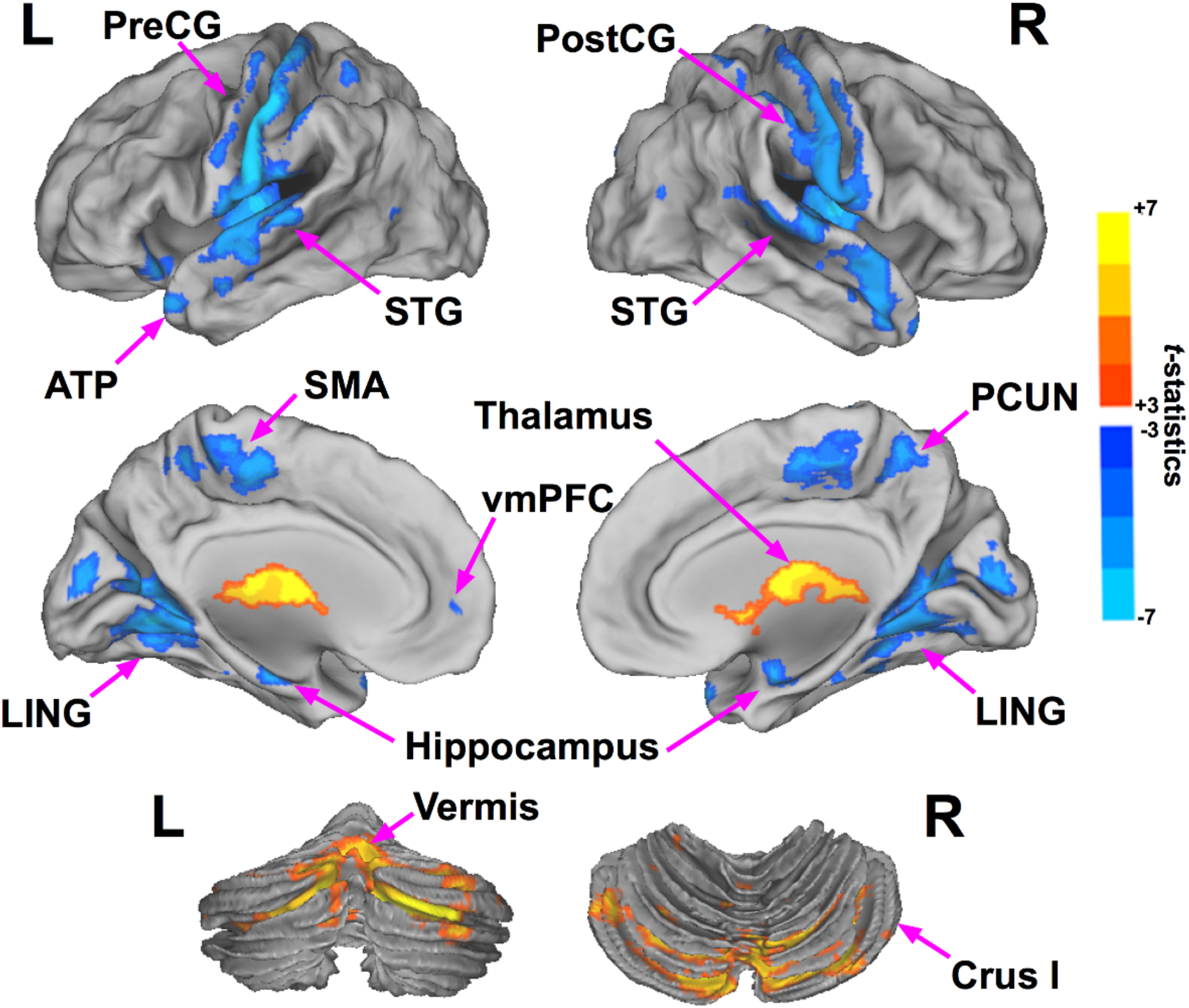
Altered functional connectivity in patients with schizophrenia when compared to healthy controls. The *t*-statistics of the group comparisons for the seed-based functional connectivity maps are shown. Red-yellow colors represent positive *t*-statistics, which indicate hypo-connectivity in the schizophrenia (SZ) group when compared to the healthy control (HC) group. Light-blue colors represent negative *t*-statistics, which indicate hyper-connectivity in the SZ group compared to the HC group. Multiple comparisons were corrected based on the false-discovery rate correction (significance level: *p* < 0.01, corrected), and an extent threshold of 20 voxels was further applied. R: right, L: left, ATP: anterior temporal pole, LING: lingual gyrus, SMA: supplementary motor area, PostCG: postcentral gyrus, PreCG: precentral gyrus, PCUN: precuneus, STG: superior temporal gyrus, and vmPFC: ventromedial prefrontal cortex.

Finally, we aimed to identify specific FCs that were associated with the experience of AVH within regions exhibiting significantly altered FC with the right crus I. Multiple linear regression analysis with PANSS (H) score as an independent variable revealed that altered FC with the left precuneus (PCUN; peak MNI coordinate = [-6, +40, +58]) exhibited highly significant positive correlations with the PANSS (H) score (Figure 5).

**Figure 5:**
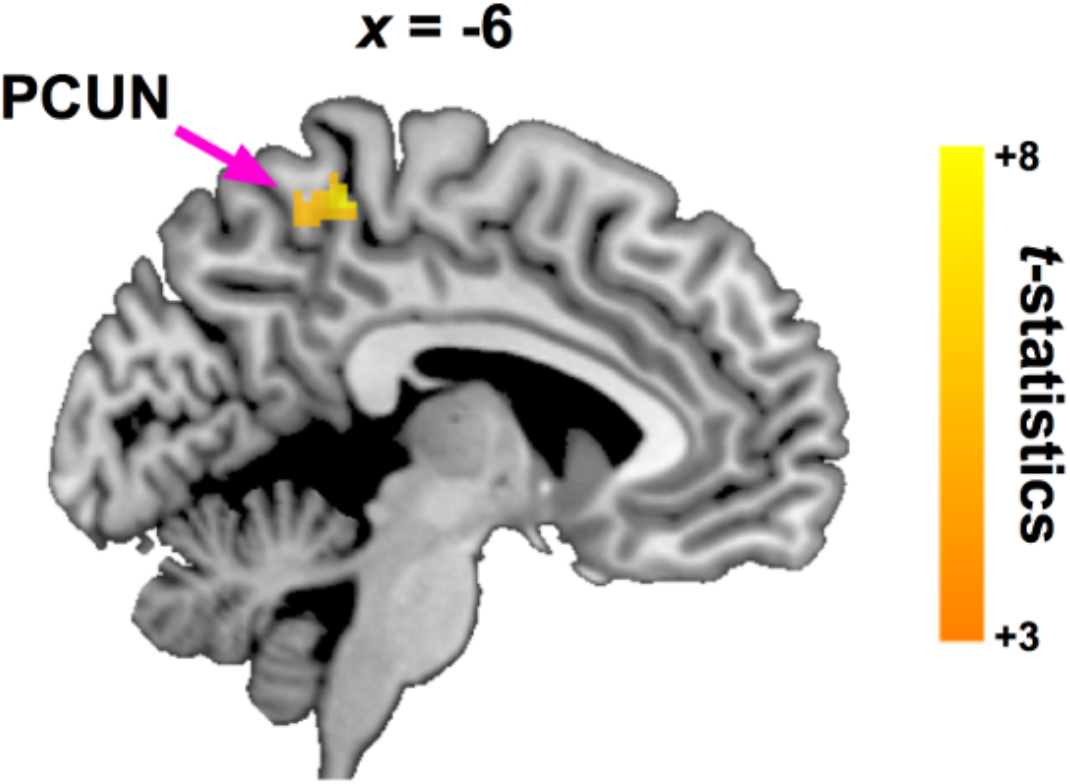
Associations between connectivity strength and the severity of auditory verbal hallucinations. Multiple linear regression analyses were performed to identify which brain regions contribute to the associations between altered principal component score in the right crus I and the severity of auditory verbal hallucinations in patients with schizophrenia. The statistical threshold was set to *p* < 0.001. Extent threshold was set to *k* = 47 voxels to achieve a family-wise error rate of *p* < 0.05 at the cluster level. Multiple linear regression analyses revealed significant positive associations between the PANSS (H) score and connectivity strength in the left precuneus (PCUN; peak MNI coordinate = [-6, +40, +58], cluster size = 55 voxels). Of note, age, handedness, gender, and the mean frame-wise displacement were included as nuisance covariates. MNI: Montreal Neurological Institute, and PANSS: Positive and Negative Syndrome Scale.

## 4. Discussion

In this study, we examined abnormal FC patterns in patients with SZ using MVPA of FC (Whitfield-Gabrieli et al., 2015) followed by post hoc seed-based FC analysis. This MVPA approach identified multiple brain regions with significant between-group differences in PC scores. Among these regions, only the right crus I had a significant association between the PC score and the PANSS (H) score. This suggests that there are alterations in some FCs involving this region that are associated with the experience of AVH in SZ. Post-hoc seed-based analysis with the right crus I as the seed indeed identified significant FC alterations such that patients with SZ exhibited hyper-connectivity in auditory, sensorimotor, and default-mode networks as well as hypo-connectivity with the cerebellar and sub-cortical networks. In particular, the strength of FC between the left PCUN and right crus I was shown to be significantly associated with the PANSS (H) score by multiple linear regression analysis. These findings suggest that, among functional abnormalities in multiple functional systems (e.g., auditory, sensorimotor and default mode networks), FC between the right crus I and left PCUN might play a critical role in the experience of AVH in patients with SZ.

The MVPA of FC enabled us to identify FC alterations associated with SZ in several clusters, such as the left STG, right SMA, bilateral thalamus, and crus I. The results were consistent with many previous neuroimaging studies reporting functional and morphological abnormalities in these regions. For instance, abnormalities in the left STG have been found at multiple levels of organization (Javitt and Sweet, 2015). These include morphological abnormalities (Hirayasu et al., 1998; Honea et al., 2008; Kasai et al., 2003), disrupted FC (Tu et al., 2015; Wang et al., 2015), and abnormal levels of activation during auditory tasks (Gaebler et al., 2015). Moreover, thalamic dysfunction, which might underlie a broad range of cognitive and behavioral impairments, has also been frequently reported (Cetin et al., 2014; Hazlett et al., 2004). Functional alterations in the thalamus showed clear laterality differences such that voxels showing increase and decrease in PC scores were found in the right and left thalamus, respectively, in the SZ group. According to a previous connectivity-based parcellation study (Behrens et al., 2003), the voxel cluster that exhibited decreased PC scores within the left thalamus were anatomically connected with temporal cortex, while the cluster that exhibited increased PC scores in the right thalamus were anatomically connected with the temporal cortex as well as frontal and posterior parietal cortices. The region-dependent FC alterations in the thalamus might reflect differences in targets of thalamic connectivity..

It is notable that a significant association between the clinical severity of AVH [i.e., PANSS (H) score] and the PC score in the right crus I of the cerebellum was found in the above clusters. The cerebellar dysfunctions have been associated with a broad range of schizophrenia symptoms, as suggested by previous studies reporting structural and functional abnormalities in the cerebellum of schizophrenia (Bernard and Mittal, 2014; Kim et al., 2014; Picard et al., 2008; Rasser et al., 2010). However, specific associations between cerebellar sub-regions and schizophrenic symptoms remained unclear. The right crus I is located between the horizontal and superior posterior fissures in the right lateral hemisphere. This cerebellar region has been shown to be critically involved in language processing (Stoodley and Schmahmann, 2009) and speech generation in the neurotypical population (Schlosser et al., 1998), whose dysfunction may contribute to the experience of AVH (Moseley et al., 2013). Consistent with our observation, resting-state functional abnormalities of this cerebellar region have been previously shown in first-episode, drug-naïve schizophrenia patients that exhibited psychotic symptoms (Guo et al., 2014; Wang et al., 2016). Therefore, aberrant connections with this area might play a vital role in the development of psychotic symptoms including AVH. Consistent with this view, post-hoc FC analysis with the right crus I as the seed revealed that patients with SZ showed hyper-connectivity with brain regions involved in auditory and sensorimotor networks as well as hypo-connectivity with the cerebellar and sub-cortical networks. Our findings indicate that AVH might stem from alterations in connections between the right crus I and regions of these functional networks.

Within these disrupted functional networks, FC between the right crus I and left PCUN exhibited a significant positive association with the PANSS (H) score. The PCUN is a brain region involved in the posterior part of the default-mode network and is essential for self-referential processing (Cavanna and Trimble, 2006). A recent meta-analytic study further suggests that this region exhibits stronger activation during other-referential processing than self-referential processing (Denny et al., 2012). Considering these previous findings, we suggest that altered FC with the PCUN may be related to aspects of AVH related to altered sense of self and other (Alderson-Day et al., 2015).

A caveat should be mentioned regarding the present methodology. Although our MVPA approach focused only on the 1st PC, it remains unknown whether and how analyses of other PCs would reveal crucial mechanisms related to AVH and other symptoms of SZ. Because the 1st PC captures the largest variance in a given dataset, we assumed that the most crucial portion of the inter-subject variations in connectivity patterns was captured by the 1st PC. Although our analysis, which focused on the primary PC, revealed associations between specific FCs and AVH, future studies need to examine the possibility that other PC scores, including the second and third PC scores, might capture different aspects of the symptomatology of SZ.

In conclusion, we adopted a combination of MVPA of FC and post-hoc FC analyses to identify FC alterations in patients with SZ without prior hypotheses regarding the seed regions. Our analyses revealed a crucial role of FC abnormalities of the right crus I with regions in auditory and sensorimotor, and default-mode networks. In particular, abnormal FC between the right crus I and left PCUN may play a vital role in the experience of AVH in SZ. Our results obtained by unbiased exploratory analysis of RSFC data highlighted the relevance of inter-regional FCs involving the cerebellum to neural abnormalities underlying SZ and AVH. These FC abnormalities might not have been identified using previous standard analyses.

## Acknowledgements

This research was supported by the Brain Mapping by Integrated Neurotechnologies for Disease Studies (Brain/MINDS) from the Japan Agency for Medical Research and Development (AMED).

## References

Alderson-Day, B., McCarthy-Jones, S., Fernyhough, C., 2015. Hearing voices in the resting brain: A review of intrinsic functional connectivity research on auditory verbal hallucinations. Neuroscience and biobehavioral reviews 55, 78–87.

Alonso-Solis, A., Vives-Gilabert, Y., Portella, M.J., Rabella, M., Grasa, E.M., Roldan, A., Keymer-Gausset, A., Molins, C., Nunez-Marin, F., Gomez-Anson, B., Alvarez, E., Corripio, I., 2017. Altered amplitude of low frequency fluctuations in schizophrenia patients with persistent auditory verbal hallucinations. Schizophrenia research.

Amad, A., Cachia, A., Gorwood, P., Pins, D., Delmaire, C., Rolland, B., Mondino, M., Thomas, P., Jardri, R., 2014. The multimodal connectivity of the hippocampal complex in auditory and visual hallucinations. Molecular psychiatry 19(2), 184–191.

Amad, A., Seidman, J., Draper, S.B., Bruchhage, M.M.K., Lowry, R.G., Wheeler, J., Robertson, A., Williams, S.C.R., Smith, M.S., 2017. Motor Learning Induces Plasticity in the Resting Brain-Drumming Up a Connection. Cerebral cortex (New York, N.Y.: 1991) 27(3), 2010–2021.

Andreasen, N.C., Pierson, R., 2008. The role of the cerebellum in schizophrenia. Biological psychiatry 64(2), 81–88.

Behrens, T.E., Johansen-Berg, H., Woolrich, M.W., Smith, S.M., Wheeler-Kingshott, C.A., Boulby, P.A., Barker, G.J., Sillery, E.L., Sheehan, K., Ciccarelli, O., Thompson, A.J., Brady, J.M., Matthews, P.M., 2003. Non-invasive mapping of connections between human thalamus and cortex using diffusion imaging. Nature neuroscience 6(7), 750–757.

Berman, R.A., Gotts, S.J., McAdams, H.M., Greenstein, D., Lalonde, F., Clasen, L., Watsky, R.E., Shora, L., Ordonez, A.E., Raznahan, A., Martin, A., Gogtay, N., Rapoport, J., 2016. Disrupted sensorimotor and social–cognitive networks underlie symptoms in childhood-onset schizophrenia. Brain: a journal of neurology 139(1), 276–291.

Bernard, J.A., Mittal, V.A., 2014. Cerebellar-motor dysfunction in schizophrenia and psychosis-risk: the importance of regional cerebellar analysis approaches. Frontiers in psychiatry 5, 160.

Cavanna, A.E., Trimble, M.R., 2006. The precuneus: a review of its functional anatomy and behavioural correlates. Brain: a journal of neurology 129(Pt 3), 564–583.

Cetin, M.S., Christensen, F., Abbott, C.C., Stephen, J.M., Mayer, A.R., Canive, J.M., Bustillo, J.R., Pearlson, G.D., Calhoun, V.D., 2014. Thalamus and posterior temporal lobe show greater internetwork connectivity at rest and across sensory paradigms in schizophrenia. NeuroImage 97, 117–126.

Cox, R.W., 1996. AFNI: software for analysis and visualization of functional magnetic resonance neuroimages. Computers and biomedical research, an international journal 29(3), 162–173.

Cui, L.B., Liu, L., Guo, F., Chen, Y.C., Chen, G., Xi, M., Qin, W., Sun, J.B., Li, C., Xi, Y.B., Wang, H.N., Yin, H., 2017. Disturbed Brain Activity in Resting-State Networks of Patients with First-Episode Schizophrenia with Auditory Verbal Hallucinations: A Cross-sectional Functional MR Imaging Study. Radiology, 160938.

de la Iglesia-Vaya, M., Escarti, M.J., Molina-Mateo, J., Marti-Bonmati, L., Gadea, M., Castellanos, F.X., Aguilar Garcia-Iturrospe, E.J., Robles, M., Biswal, B.B., Sanjuan, J., 2014. Abnormal synchrony and effective connectivity in patients with schizophrenia and auditory hallucinations. NeuroImage. Clinical 6, 171–179.

Denny, B.T., Kober, H., Wager, T.D., Ochsner, K.N., 2012. A meta-analysis of functional neuroimaging studies of self- and other judgments reveals a spatial gradient for mentalizing in medial prefrontal cortex. Journal of cognitive neuroscience 24(8), 1742–1752.

Gaebler, A.J., Mathiak, K., Koten, J.W., Jr., Konig, A.A., Koush, Y., Weyer, D., Depner, C., Matentzoglu, S., Edgar, J.C., Willmes, K., Zvyagintsev, M., 2015. Auditory mismatch impairments are characterized by core neural dysfunctions in schizophrenia. Brain: a journal of neurology 138(Pt 5), 1410–1423.

Guo, W., Yao, D., Jiang, J., Su, Q., Zhang, Z., Zhang, J., Yu, L., Xiao, C., 2014. Abnormal defaultmode network homogeneity in first-episode, drug-naive schizophrenia at rest. Prog Neuropsychopharmacol Biol Psychiatry 49, 16–20.

Hazlett, E.A., Buchsbaum, M.S., Kemether, E., Bloom, R., Platholi, J., Brickman, A.M., Shihabuddin, L., Tang, C., Byne, W., 2004. Abnormal glucose metabolism in the mediodorsal nucleus of the thalamus in schizophrenia. The American journal of psychiatry 161(2), 305–314.

Hirayasu, Y., Shenton, M.E., Salisbury, D.F., Dickey, C.C., Fischer, I.A., Mazzoni, P., Kisler, T., Arakaki, H., Kwon, J.S., Anderson, J.E., Yurgelun-Todd, D., Tohen, M., McCarley, R.W., 1998. Lower left temporal lobe MRI volumes in patients with first-episode schizophrenia compared with psychotic patients with first-episode affective disorder and normal subjects. The American journal of psychiatry 155(10), 1384–1391.

Hoffman, R.E., Hawkins, K.A., Gueorguieva, R., Boutros, N.N., Rachid, F., Carroll, K., Krystal, J.H., 2003. Transcranial magnetic stimulation of left temporoparietal cortex and medication-resistant auditory hallucinations. Archives of general psychiatry 60(1), 49–56.

Honea, R.A., Meyer-Lindenberg, A., Hobbs, K.B., Pezawas, L., Mattay, V.S., Egan, M.F., Verchinski, B., Passingham, R.E., Weinberger, D.R., Callicott, J.H., 2008. Is gray matter volume an intermediate phenotype for schizophrenia? A voxel-based morphometry study of patients with schizophrenia and their healthy siblings. Biological psychiatry 63(5), 465–474.

Jardri, R., Thomas, P., Delmaire, C., Delion, P., Pins, D., 2013. The neurodynamic organization of modality-dependent hallucinations. Cerebral cortex (New York, N.Y.: 1991) 23(5), 1108–1117.

Javitt, D.C., Sweet, R.A., 2015. Auditory dysfunction in schizophrenia: integrating clinical and basic features. Nature reviews. Neuroscience 16(9), 535–550.

Kasai, K., Shenton, M.E., Salisbury, D.F., Hirayasu, Y., Lee, C.U., Ciszewski, A.A., Yurgelun-Todd, D., Kikinis, R., Jolesz, F.A., McCarley, R.W., 2003. Progressive decrease of left superior temporal gyrus gray matter volume in patients with first-episode schizophrenia. The American journal of psychiatry 160(1), 156–164.

Kim, D.J., Kent, J.S., Bolbecker, A.R., Sporns, O., Cheng, H., Newman, S.D., Puce, A., O’Donnell, B.F., Hetrick, W.P., 2014. Disrupted modular architecture of cerebellum in schizophrenia: a graph theoretic analysis. Schizophrenia bulletin 40(6), 1216–1226.

Matsuoka, K., Uno, M., Kasai, K., Koyama, K., Kim, Y., 2006. Estimation of premorbid IQ in individuals with Alzheimer’s disease using Japanese ideographic script (Kanji) compound words: Japanese version of National Adult Reading Test. Psychiatry and clinical neurosciences 60(3), 332–339.

McKeown, M.J., Hansen, L.K., Sejnowsk, T.J., 2003. Independent component analysis of functional MRI: what is signal and what is noise? Curr Opin Neurobiol 13(5), 620–629.

Moseley, P., Fernyhough, C., Ellison, A., 2013. Auditory verbal hallucinations as atypical inner speech monitoring, and the potential of neurostimulation as a treatment option. Neuroscience and biobehavioral reviews 37(10 Pt 2), 2794–2805.

Murphy, K., Birn, R.M., Handwerker, D.A., Jones, T.B., Bandettini, P.A., 2009. The impact of global signal regression on resting state correlations: are anti-correlated networks introduced? NeuroImage 44(3), 893–905.

Oldfield, R.C., 1971. The assessment and analysis of handedness: the Edinburgh inventory. Neuropsychologia 9(1), 97–113.

Picard, H., Amado, I., Mouchet-Mages, S., Olie, J.P., Krebs, M.O., 2008. The role of the cerebellum in schizophrenia: an update of clinical, cognitive, and functional evidences. Schizophrenia bulletin 34(1), 155–172.

Power, J.D., Barnes, K.A., Snyder, A.Z., Schlaggar, B.L., Petersen, S.E., 2012. Spurious but systematic correlations in functional connectivity MRI networks arise from subject motion. NeuroImage 59(3), 2142–2154.

Rasser, P.E., Schall, U., Peck, G., Cohen, M., Johnston, P., Khoo, K., Carr, V.J., Ward, P.B., Thompson, P.M., 2010. Cerebellar grey matter deficits in first-episode schizophrenia mapped using cortical pattern matching. NeuroImage 53(4), 1175–1180.

Rolland, B., Amad, A., Poulet, E., Bordet, R., Vignaud, A., Bation, R., Delmaire, C., Thomas, P., Cottencin, O., Jardri, R., 2015. Resting-state functional connectivity of the nucleus accumbens in auditory and visual hallucinations in schizophrenia. Schizophrenia bulletin 41(1), 291–299.

Saad, Z.S., Gotts, S.J., Murphy, K., Chen, G., Jo, H.J., Martin, A., Cox, R.W., 2012. Trouble at rest: how correlation patterns and group differences become distorted after global signal regression. Brain connectivity 2(1), 25–32.

Schlosser, R., Hutchinson, M., Joseffer, S., Rusinek, H., Saarimaki, A., Stevenson, J., Dewey, S.L., Brodie, J.D., 1998. Functional magnetic resonance imaging of human brain activity in a verbal fluency task. Journal of neurology, neurosurgery, and psychiatry 64(4), 492–498.

Shergill, S.S., Brammer, M.J., Williams, S.C., Murray, R.M., McGuire, P.K., 2000. Mapping auditory hallucinations in schizophrenia using functional magnetic resonance imaging. Archives of general psychiatry 57(11), 1033–1038.

Sommer, I.E., Clos, M., Meijering, A.L., Diederen, K.M., Eickhoff, S.B., 2012. Resting state functional connectivity in patients with chronic hallucinations. PloS one 7(9), e43516.

Stoodley, C.J., Schmahmann, J.D., 2009. Functional topography in the human cerebellum: A metaanalysis of neuroimaging studies. NeuroImage 44(2), 489–501.

Thoma, R.J., Chaze, C., Lewine, J.D., Calhoun, V.D., Clark, V.P., Bustillo, J., Houck, J., Ford, J., Bigelow, R., Wilhelmi, C., Stephen, J.M., Turner, J.A., 2016. Functional MRI Evaluation of Multiple Neural Networks Underlying Auditory Verbal Hallucinations in Schizophrenia Spectrum Disorders. Frontiers in psychiatry 7, 39.

Tu, P.C., Lee, Y.C., Chen, Y.S., Hsu, J.W., Li, C.T., Su, T.P., 2015. Network-specific cortico-thalamic dysconnection in schizophrenia revealed by intrinsic functional connectivity analyses. Schizophrenia research 166(1-3), 137–143.

Wang, H., Guo, W., Liu, F., Wang, G., Lyu, H., Wu, R., Chen, J., Wang, S., Li, L., Zhao, J., 2016. Patients with first-episode, drug-naive schizophrenia and subjects at ultra-high risk of psychosis shared increased cerebellar-default mode network connectivity at rest. Scientific reports 6, 26124.

Wang, H.L., Rau, C.L., Li, Y.M., Chen, Y.P., Yu, R., 2015. Disrupted thalamic resting-state functional networks in schizophrenia. Frontiers in behavioral neuroscience 9, 45.

Whitfield-Gabrieli, S., Ghosh, S.S., Nieto-Castanon, A., Saygin, Z., Doehrmann, O., Chai, X.J., Reynolds, G.O., Hofmann, S.G., Pollack, M.H., Gabrieli, J.D., 2015. Brain connectomics predict response to treatment in social anxiety disorder. Molecular psychiatry.

Yang, G.J., Murray, J.D., Glasser, M., Pearlson, G.D., Krystal, J.H., Schleifer, C., Repovs, G., Anticevic, A., 2016. Altered Global Signal Topography in Schizophrenia. Cerebral cortex (New York, N.Y.: 1991).

Yang, G.J., Murray, J.D., Repovs, G., Cole, M.W., Savic, A., Glasser, M.F., Pittenger, C., Krystal, J.H., Wang, X.J., Pearlson, G.D., Glahn, D.C., Anticevic, A., 2014. Altered global brain signal in schizophrenia. Proceedings of the National Academy of Sciences of the United States of America 111(20), 7438–7443.

Zhu, J., Wang, C., Liu, F., Qin, W., Li, J., Zhuo, C., 2016. Alterations of Functional and Structural Networks in Schizophrenia Patients with Auditory Verbal Hallucinations. Frontiers in human neuroscience 10, 114.

